# Autoreactive B cells in rheumatoid arthritis consist of activated CXCR3+ memory B cells and plasmablasts

**DOI:** 10.1101/2023.05.19.538699

**Authors:** Sanne Reijm, Joanneke C. Kwekkeboom, Nienke J. Blomberg, Jolien Suurmond, Diane van der Woude, Rene E.M. Toes, Hans U. Scherer

## Abstract

Many autoimmune diseases (AIDs) are characterized by persistence of autoreactive B cell responses which is often directly implicated in disease pathogenesis. How and why these cells are generated or how they are maintained for years is largely unknown. Rheumatoid arthritis is among the most common AIDs and characterized by autoantibodies recognizing proteins with post-translational modifications (PTMs). This PTM-directed, autoreactive B cell compartment is ill defined. Here, we visualized the B cell response against the three main types of PTM antigens implicated in RA by spectral flow cytometry. Our results show extensive cross-reactivity of autoreactive B cells against all three PTM antigens (citrulline, homocitrulline and acetyllysine). Unsupervised clustering revealed several distinct memory B cell (mBC) populations. Autoreactive cells clustered with the most recently activated, class-switched mBC phenotype, expressing high CD80, low CD24 and low CD21. Notably, patients also harbored large fractions of autoreactive plasmablasts (PB). Both PTM-directed mBC and PB showed high expression of CXCR3, a receptor for chemokines abundantly present in arthritic joints. Together, our data provide novel, detailed insight into the biology of B cell autoreactivity and its remarkable, seemingly exhaustless persistence in a prominent human AID.

## Introduction

Many autoimmune diseases (AIDs), such as rheumatoid arthritis (RA), pemphigus vulgaris, systemic lupus erythematosus or myasthenia gravis, are characterized by the presence of disease-specific autoantibodies. Their presence and the remarkable efficacy of B cell depletion indicate a central role for autoreactive B cells in disease pathogenesis (1-4). While autoantibody responses are often well characterized, little information is present on the phenotype and composition of the underlying, autoreactive B cell compartments. Autoreactive B cell memory frequently persists for a lifetime and is difficult to eliminate therapeutically. Such lifetime memory would be desirable for vaccine responses but information about the differences between autoreactive B cells in comparison to B cells induced by recall antigens or infections is sparse. Continuous and often systemic presence of antigens that immune cells can recognize is an hallmark of autoimmunity, while the presence of recall antigens is frequently transient and localized. Such different dynamics in antigen-exposure can affect the outcome of immune responses. T cells exposed to immunogenic tumors, for example, lose responsiveness by expressing inhibitory molecules and/or exhaustion markers such as PD-1 and CTLA-4. Vaccine responses, on the other hand, generate memory response that can persist for life (5). The dynamics of memory B cell responses are, in this context, less well defined. One could, for example, expect signs of exhaustion or decay of the autoreactive memory B cell compartment in time, especially in case the autoantigen is continuously and systemically present. The observation that circulating plasma cells can differentiate from memory B cells (mBC) is, therefore, intriguing in an autoimmune context, as it may contain valuable clues as to how such responses can be maintained (6). In HIV infection, atypical B cells, also called double-negative 2 (DN2) or age-associated B cells, are IgD^-^CD27^-^CD21^-^ B cells which have been described to be dysfunctional or exhausted, while in SLE these B cells are linked to the production of autoantibodies (7, 8). Also in RA, atypical B cells are associated with disease, but it is unknown if these cells are autoreactive (9). To better comprehend the biology of antigen-specific, autoreactive B cell populations in the human, we embarked on the visualization of autoreactive B cells recognizing multiple autoantigens in RA, a prototypic AID with immunological features (such as HLA association and autoantibodies) inherent to other, less-frequent yet equally debilitating B cell driven AIDs.

RA is characterized by systemic inflammation and a localized, destructive inflammatory process in the joints. The presence of anti-modified protein antibodies (AMPA) hallmarks the disease and is found in approximately 70% of patients. AMPA can contain reactivities to different post-translational modifications (PTMs), including anti-citrullinated protein antibodies (ACPA), anti-carbamylated protein antibodies (anti-CarP) and anti-acetylated protein antibodies (AAPA). How AMPA responses are generated is currently unknown. Secreted AMPA can be highly cross-reactive to citrullinated, carbamylated and/or acetylated antigens with varying affinities (10-12). This cross-reactivity presumably impacts on B cell activation in vivo, as human B cell lines expressing B cell receptors (BCR) recognizing citrullinated antigens can be cross-activated by carbamylated or acetylated proteins (13). The extent of cross-reactivity at the (memory) B cell level is still ill-defined, however, as cross-reactivity has so far been analyzed primarily for AMPA-responses in serum and for monoclonal ACPA. It is, for example, conceivable that the secreted AMPA responses are derived from small fractions of cross-reactive B cells as these could, potentially, become most activated. Insights into the cross-reactivity of autoreactive B cells would, therefore, be valuable as it could help to define the nature of PTMs to which AMPA-expressing memory B cells are predominantly directed and which stimulate their activation.

Recently, we identified citrullinated protein (Cit)-directed mBCs in RA and revealed that these cells express an activated phenotype based on the expression of CD80, CD86, HLA-DR and Ki-67 (14). Exploiting the possibilities of spectral flowcytometry, we now generated a combinatorial staining approach to analyze the autoreactive PB-compartment next to an extensive phenotypic evaluation of the autoreactive memory population to multiple PTM-antigens. Our results show the presence of a considerable population of autoreactive PB and point to the continuous differentiation of (activated), cross-reactive mBCs towards PB in chronic disease. This persistent activity of the autoreactive B cell compartment may be involved in the chronicity of RA, a disease that frequently flares if anti-inflammatory treatment is stopped. Hence, extensive phenotyping and improved staining methods reveal insights in autoreactive B cell differentiation in RA, with possible implications for autoreactive B cell targeting, tolerance induction but also efforts in the vaccine field aiming at generating long-lasting B cell memory.

## Results

### Citrullinated antigen-directed B cells are most abundant within the PTM-directed B cell population

We first set out to define the (cross)reactivity of the autoreactive B cell population based on the different PTM-reactivities associated with RA. We visualized the different ‘classes’ of autoantigen-specific B cells (Citrullinated protein (Cit)-, Carbamylated protein (CarP)-, Acetylated protein(Acetyl)-directed B cells) in RA patients by using differentially labelled streptavidin-tetramers carrying individual PTM-modified peptides. Two tetramers with different fluorescent labels were generated for each modification, together with tetramers carrying control peptides with the non-modified backbone amino acids lysine and arginine. Cells were stained simultaneously with all tetramers. This approach allowed us to obtain an antigen-specific double staining signal per reactivity. To optimize the staining, we applied human Ramos B cell lines transduced with BCRs recognizing the different PTM-antigens with pre-determined cross-reactivity profiles (13). Ramos-2G9 cells stained positive for CCP4 and CHCitP4 tetramers in this setting, for example, while BCR-deficient Ramos cells (Ramos-MDL) did not show binding (Figure S1). Likewise, the acetylated antigen-directed cell line Ramos-7E4 stained positive for CAcetylP4 tetramers. Subsequently, a panel was designed to visualize and phenotype all PTM-directed B cells. Labelled antigens to identify TT-directed B cells and a set of markers associated with B cell activation and homing were additionally included (table S3). Staining B cells enriched from PBMCs from RA patients revealed that Cit-, CarP-, Acetyl-, and TT-directed B cells could all be detected within the same patient and within one sample (Figure 1A). The specificity of the Cit staining was confirmed in previous studies (15) and the CarP- and Acetyl-directed B cell staining was further confirmed in inhibition experiments, using tetramers without fluorescent label which were added to the cells in excess before staining with the labeled tetramers (Figure S2). TT-directed cells were detected in similar frequencies in RA patients and HD (Figure 1B). On the other hand, PTM-directed B cells could be detected almost exclusively in RA patients, further confirming the specificity of the autoreactive B cell staining (Figure 1B). Among the PTM-directed B cells, Cit-directed B cells were most frequent, followed by CarP-directed B cells. Acetyl-directed B cells could only be detected in a minority of RA patients. Finally, few Acetyl-directed B cells could also be detected in HD. Together, these data show the feasibility to simultaneously visualize 3 different autoreactive B cell populations and a comparator B cell population against a recall antigen in samples of individual patients.

**Figure 1.**
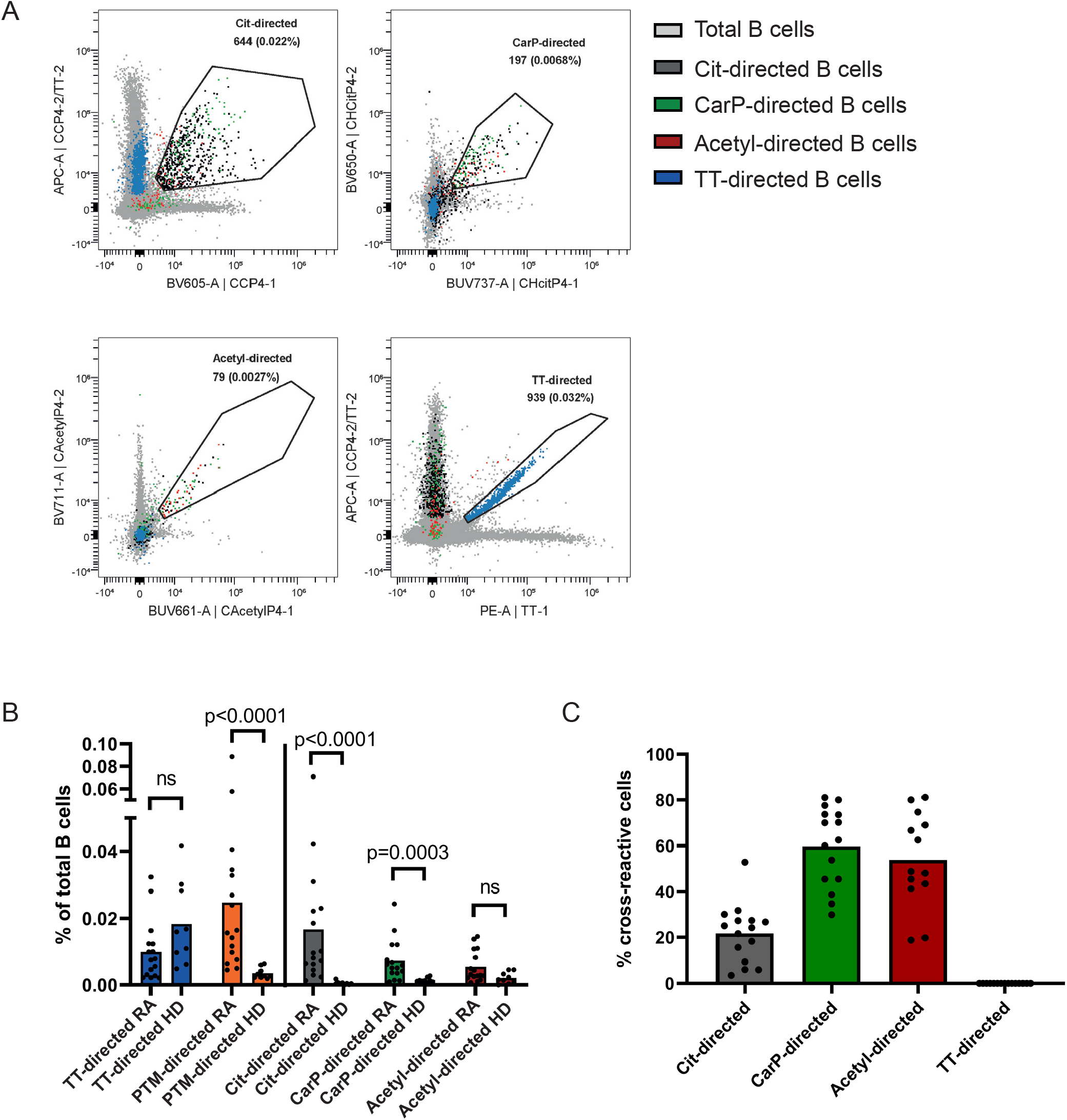
A) Identification of Cit-, CarP-, Acetyl- and TT-directed B cells. Example of 1 RA patient. B) Quantification of PTM- and TT-directed B cells in percent of total B cells in RA patients (n=16) and HD (n=9). Each dot represents one individual, bars represent means. Statistical testing performed with Mann-Whitney U test. C) Cross-reactivity of PTM- and TT-directed B cells in RA patients to at least one other PTM (n=16). Each dot represents one individual, bars represent means.

### PTM-directed B cells are highly cross-reactive with citrulline as dominant antigen

Cross-reactivity within (autoreactive) B cell responses is an intriguing phenomenon, as it may point to (auto-)antigens driving B cell activation beyond commonly anticipated routes. It has previously been shown that AMPA in serum are highly cross-reactive towards different PTM-antigens. To understand the degree of cross-reactivity on the level of circulating B cells, we next assessed cross-reactivity of the B cell receptors. Our results show that a large fraction of PTM-directed B cells in the circulation react to multiple modified antigens, indicating that these cells are, indeed, cross-reactive (Figure 1A). More specifically, around 20% of Cit-directed B cells recognized at least 1 other PTM antigen, while this was over 50% for Acetyl- and CarP-directed B cells (Figure 1C). This proportion is likely an underestimation of cross-reactive cells, as the frequency of CarP-directed B cells increased when cit- and acetyl-containing tetramers were omitted from the staining and, hence, could not compete for binding (Figure S3). These findings indicate that a substantial proportion of modified-antigen directed cells in RA is cross-reactive to other PTMs, while others show mono-reactivity. Cit-directed B cells were most frequent and more often mono-reactive than CarP- and Acetyl-directed B cells, suggesting that citrulline is the dominant antigen in the AMPA response. Furthermore, these results indicate that a combined staining for all three PTMs is required to visualize the full breadth of the PTM-directed B cell response in RA as not all cells show cross-reactivity to other PTM antigens.

### The composition of the overall B cell population in HD and RA patients

To place the PTM-directed B cell response into the context of the total B cell population and its possible alterations in RA, we next identified B cell subsets by unsupervised FlowSom clustering using normalized expression levels of the markers CD19, CD20, CD21, CD24, CD27, CD38, IgD and IgM. This resulted in 20 different B cell clusters (Figure 2A and S4). B cell subsets were assigned names based on the expression of the markers depicted in Figure 2C and S4 (16). Interestingly, 3 different CD27^+^ switched memory subsets were identified based on the differential expression of CD20, CD21 and CD24. “Switched memory-1” cells expressed intermediate levels of CD20 and high levels of CD21 and CD24. “Switched “memory-2” B cells were CD20^int^CD21^+^CD24^lo^, while “switched memory-3” B cells were CD20^hi^CD21^-^CD24^-^ (Figure 2D). Another interesting subset was found which clustered with PB (cluster 19). These cells were similar to switched memory-3 cells, except that the subset lacked CD20 expression. The cells had a low expression of CD62L, indicating PB fate, but also low expression of CD38, indicating that they were not fully differentiated PB (17). Therefore, the subset likely consists of cells differentiating towards PB and were called pre-PB. To determine differences in overall B cell distribution between HD and RA patients, the B cells from these groups were visualized as depicted in Figure 2B. Differences were quantified for the switched memory clusters, DN2 cells (CD20^+^CD27^-^IgD^-^CD21^-^) and PB clusters (figures 2E, F and G). No major differences were found between RA patients and HD in the switched memory clusters, although a small decrease for switched memory-1 was found in patients. Some RA patients showed higher percentages of DN2 cells (Figure 2F), albeit the difference was not significant. Furthermore, RA patients displayed a higher percentage of total pre-PB. Finally, 5 out of 16 RA patients displayed a high percentage (above 4% of total B cells) of switched PB while this was not found in any of the HD.

**Figure 2.**
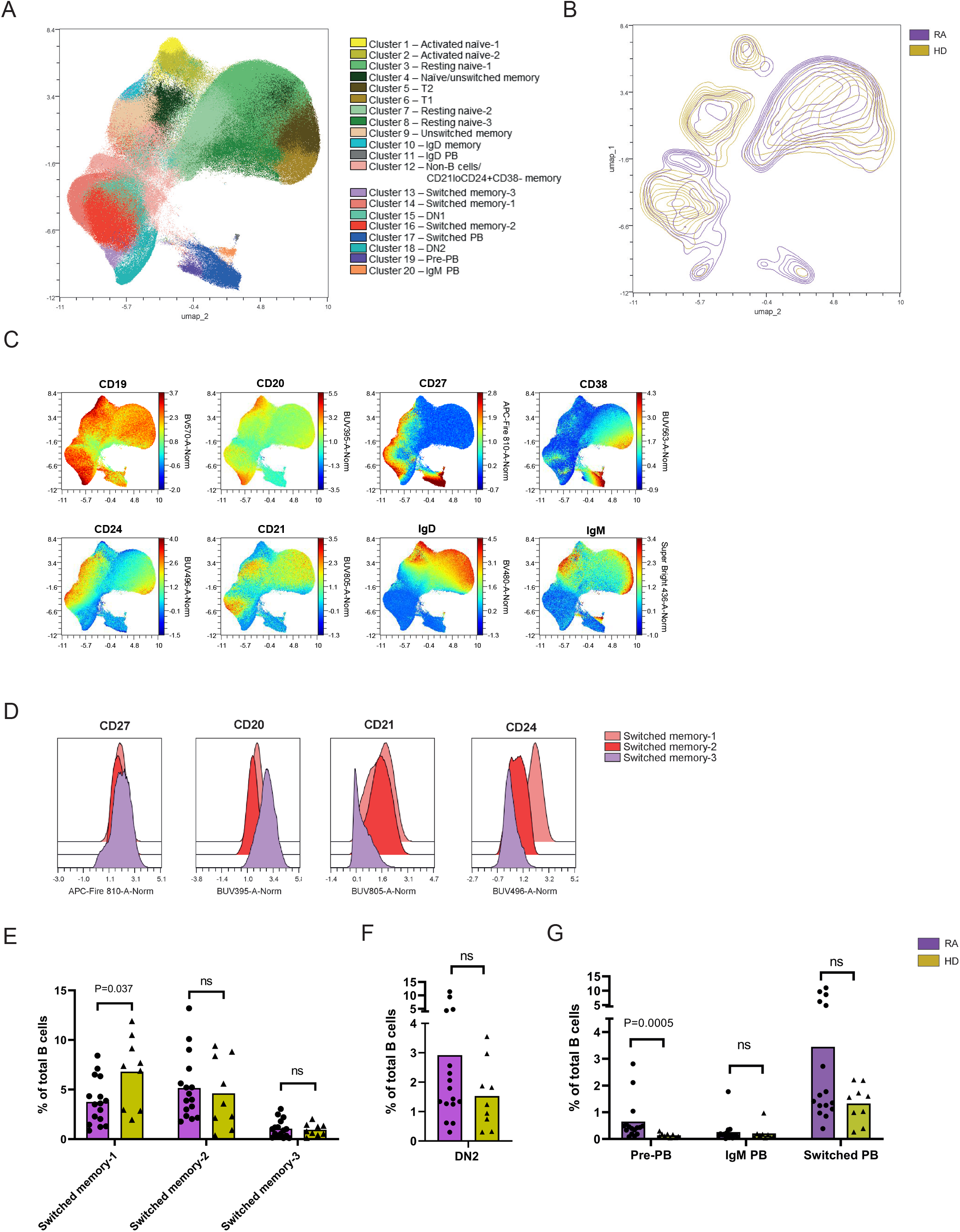
A) UMAP visualization of B cell clusters based on the markers depicted in C. B) Contour plot of 16 RA patients (purple) and 9 HD (yellow) on UMAP. C) Expression levels (in normalized MFI) of CD19, CD20, CD27, CD38, CD24, CD21, IgD and IgM. D) Histograms of CD27, CD20, CD21 and CD24 expression in switched memory-1, 2 and 3 subsets. E, F and G) Percentages of 3 switched memory B cell subsets, DN2 and 3 PB subsets of total B cells from RA patients (purple) and HD (yellow). Each dot represents one individual, bars represent means. Statistical testing was performed with Mann-Whitney U test.

### The PTM-directed B cell population is hallmarked by recently activated IgG+ memory B cells and a high frequency of PBs

To further define the phenotypes of PTM-directed B cells and their non-autoreactive TT-directed comparators in this context, the same clustering strategy was applied to these antigen-specific responses. PTM-directed B cells from RA patients clustered mainly with the populations of pre-PB, switched PB and class-switched mBC (Figure 3A). Subset analysis of Cit-, CarP and Acetyl-directed B cells revealed that these responses were remarkably similar (Figure S5A and B). Selective expansions of autoreactive PB directed against individual PTMs were not observed. The few PTM-directed B cells that were detectable in HD did not cluster to a specific compartment but were scattered throughout the plot and mainly present in the resting naïve B cell clusters (Figure 3A). TT-directed B cells, on the other hand, clustered with IgM PB and class-switched mBC. Interestingly, TT- and PTM-directed B cells clustered with different subsets of switched mBC. More specifically, the highest percentages of memory PTM-directed B cells were found within the switched memory-2 and switched memory-3 clusters, while the memory TT-directed B cells were almost exclusively present in switched memory-1 and switched memory-2 clusters (figure 3B). In the PB compartment, most PTM-directed B cells were found within the switched PB cluster, but also the pre-PB cluster contained a relatively high percentage of PTM-directed cells. Of note, no correlation was found between the percentage of PTM-directed PB and the total percentage of PB (Figure 3C). The latter suggests that the phenotype of PTM-directed B cells is not influenced by an overrepresentation of these B cell subsets in RA patients.

**Figure 3.**
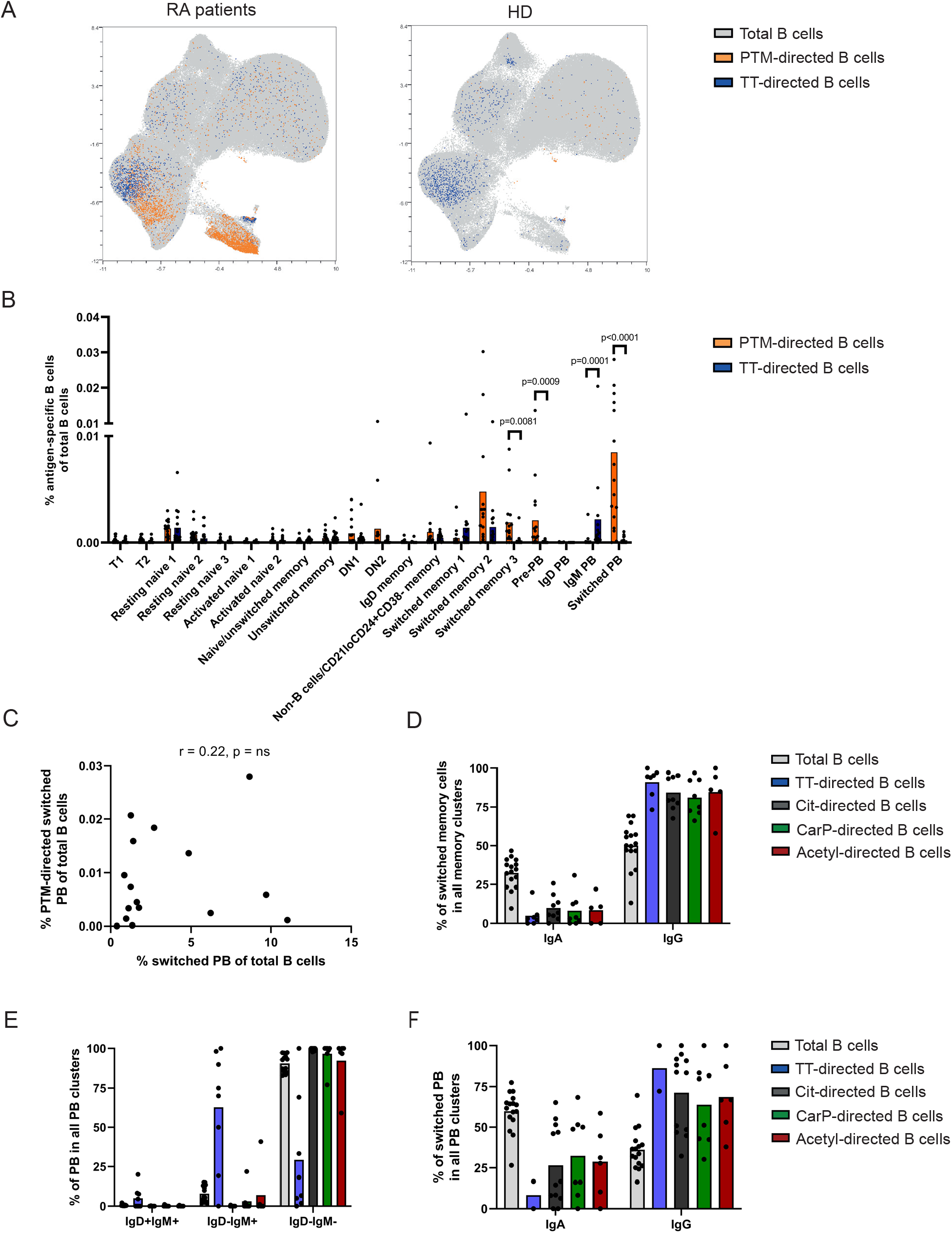
A) UMAP visualization of total B cells (grey), PTM- (orange) and TT-directed B cells (blue) of RA patients (n=16) on the left or HD (n=9) on the right. B) Percentage of PTM- and TT-directed B cells of total B cells separated by the 20 identified B cell subsets. Each dot represents one individual, bars represent means. Statistical testing was performed with Wilcoxon signed rank test. C) Correlaon between the percentage of PTM-directed switched PB and the percentage of total switched PB of total B cells. D) Percentages of IgA and IgG expression cells in the switched memory cells. E) Percentages of IgM and IgD expressing cells in the switched PB subsets. F) Percentages of IgA and IgG expressing cells in the switched PB subsets. Each dot represents one individual, bars represent means.

Class-switch recombination is influenced by the environment in which B cells mature. Therefore, isotype use within the autoreactive B cell compartment could provide clues about the site of B cell activation. IgA, for example, is mostly induced at mucosal sites under the influence of TGF-β (18). Within the compartment of IgM^-^IgD^-^ class-switched mBC, the vast majority of the PTM-directed B cells and TT-directed B cells expressed IgG BCRs (Figure 3D). The PB subsets showed a similar pattern as most PTM-directed PB were class-switched (Figure 3E). IgG^+^ cells dominated the PTM-directed PB compartment, in line with their distribution within class-switched mBC (figure 3F). Nonetheless, there was an expansion of IgA^+^ autoreactive B cells in about 50% of RA patients. In addition, the percentage of IgG or IgA B cells towards a PTM corresponded in general with plasma levels of the respective AMPAs (Figure S6). These data suggest a close link between the autoreactive mBC and the PB compartment.

### PTM-directed B cells may home to sites of inflammation

The activation of the PTM-directed B cell compartment and the presence of a large PB fraction are intriguing. PBs are migratory cells capable of ‘carrying’ local immune responses to distant tissues. As neither the site of generation and activation of PTM-directed B cells nor the mechanisms involved in the initiation of tissue inflammation in RA are well understood, we opted to characterize activation and homing marker profiles of these cells. To this end, median fluorescent intensities (MFI) of all markers were determined within the different mBC and PB clusters (Figure 4, S7 and S8). Clustering confirmed that TT-directed and PTM-directed B cells belong to different mBC subsets (figure 4A and S6). Increasing expression of CD80, CD86 and Ki-67 in switched memory 1, 2 and 3 clusters, respectively, underlined that PTM-directed mBC are among the most recently activated mBC in peripheral blood. In contrast, TT-directed B cells remained in a resting state.

**Figure 4.**
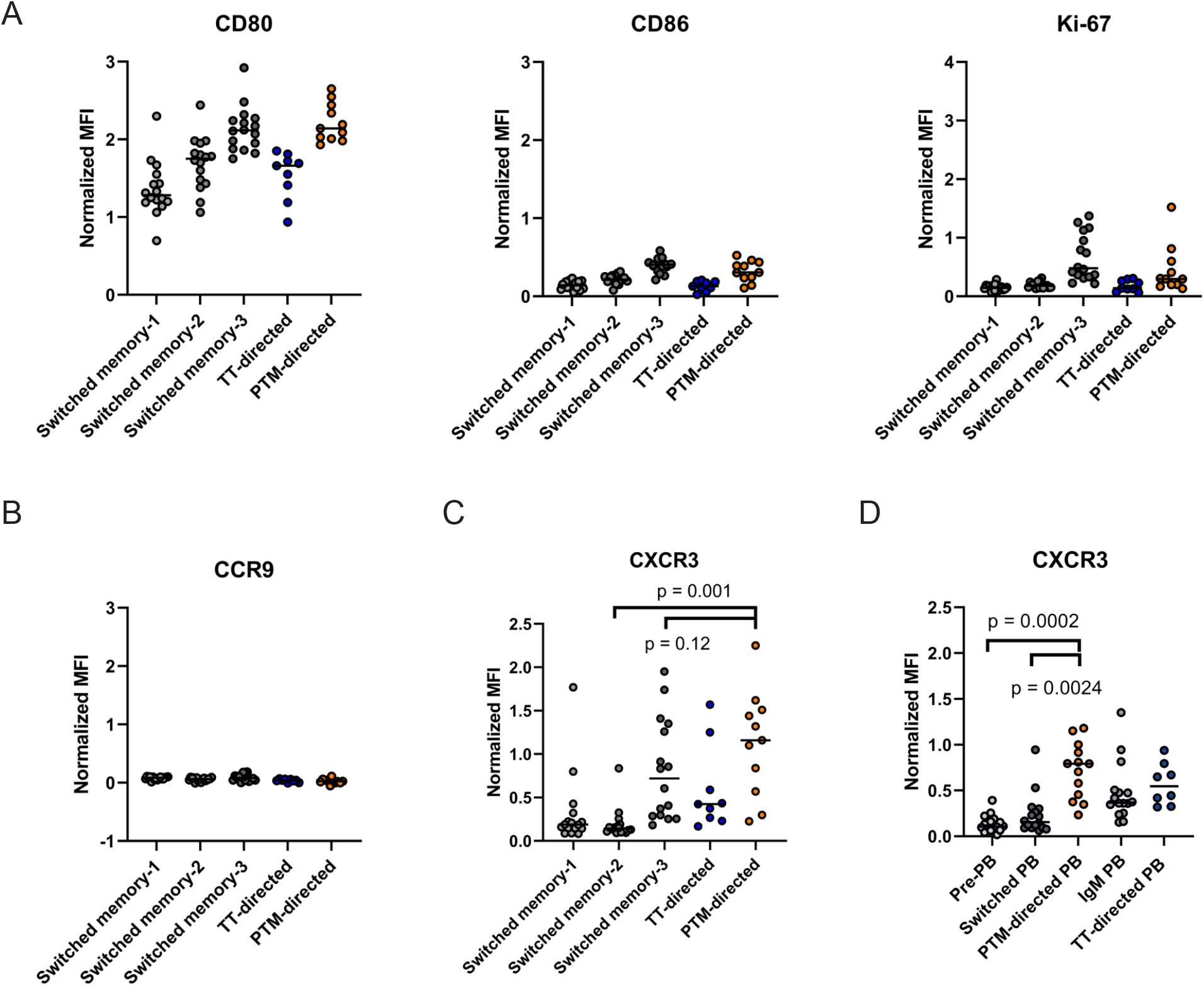
A) Normalized MFIs of CD80, CD86 and Ki-67 in 3 switched memory clusters and of TT-directed B cells and PTM-directed B cells within these clusters. B) Normalized MFI of CCR9 in 3 switched memory clusters and of TT-directed B cells and PTM-directed B cells within these clusters. C) Normalized MFI of CXCR3 in the 3 switched memory clusters and (D) in the 3 PB clusters, including TT-directed B cells and PTM-directed B cells within these clusters. Differences in CXCR3 expression were statistically tested with the Wilcoxon signed rank test. In all plots in the figure, each dot represent 1 RA patient, black lines represent medians. Only those PTM- and TT-directed B cells that were present in 1 of the 3 switched memory clusters (A-C) or 1 of the 3 PB clusters (D) were included in the analysis. Donors with less than 10 TT- or PTM-directed B cells in the analyzed clusters were excluded from the analysis.

A common hypothesis about the origin of PTM-directed B cells points to mucosal tissues. To gather indications in support of this hypothesis, the expression of the gut-homing marker CCR9 was analyzed (18). CCR9 was absent on mBC, including PTM- and TT-directed B cells (Figure 4B). IgM PB expressed some CCR9 as did IgM TT-directed PB, while PTM-directed B cells did not display this marker (Figure S8). As such, these observations suggest that PTM-directed B cells circulating in the peripheral blood do not specifically home to (or originate from) gut mucosal sites (19).

Interestingly, the switched memory-3 B cell cluster expressed high levels of the chemokines receptor CXCR3. The expression of this receptor tended to be even higher in PTM-directed memory B cells, although this difference did not reach significance (Figure 4C). In line with this finding, CXCR3 expression on PTM-directed PB was increased significantly compared to all B cells in the switched (pre-)PB subsets (Figure 4D). CXCR3 expression is correlated with IgG1 isotype expression (20). As PTM-directed B cells are mainly skewed to the IgG isotype, we compared all IgG memory cells and PB to PTM-directed IgG memory cells and PB. Interestingly, CXCR3 expression was still elevated in PTM-directed B cells (Figure S9), indicating a selective upregulation of this chemokine receptor on autoreactive B cells. CXCR3 mediates migration of B cells towards ligands present in inflamed tissues. Hence, PTM-directed B cells in the peripheral blood are likely responsive to chemokine gradients guiding them towards the inflamed synovium in RA patients.

## Discussion

Autoreactive B cells play a major role in many AIDs. In the absence of suitable murine models that closely reflect anti-PTM autoreactivity as observed in human RA, and considering the well-known differences observed between human and murine (memory) B cell populations, we here focused on elaborate phenotyping of PTM-directed B cells in human samples. We performed a study which is unique in how it addresses the complexity of the autoreactive B cell compartment, taking into account its heterogeneity as well as its remarkable cross-reactivity. Intracellular staining with isotypes and labeled antigens allowed us to visualize PB in addition to other B cell subsets. We found that ∼50% of circulating PTM-directed B cells were indeed PB. The size of this compartment is considerably larger than previously anticipated (14). This is likely a consequence of the challenges of incorporating PB into B cell panels for antigen-specific studies which requires intra-cellular staining, together with their vulnerability to cryopreservation (21). All analyses here were performed on fresh cells only, using protocols that allow for the visualization of both PB and mBCs by spectral flowcytometry. Phenotypically, PTM-directed mBC displayed a phenotype (CD19^hi^CD20^+^CD27^+^CD21^-^ CD24^-^) which is often attributed to activated memory B cells. Loss of CD21 and CD24 and high expression of costimulatory factors such as CD80 indicate recent B cell activation (16, 22). Furthermore, CD27^+^CD21^-^ mBC have been linked to cells that recently left the germinal center (23). Such cells were described to express high levels of co-stimulatory receptors and low CCR7, which is in concordance with the PTM-directed B cell phenotype observed in this study. These mBC might be progenitors of PBs, as similar clones have been found within this activated mBC subset and within PB (23, 24). Interestingly, we also found PTM-directed B cells which displayed a phenotype “in between” activated mBC and PB. The pre-PB population supports this notion. Of note, the relative frequency of auto-antigen-directed PB observed in some RA patients seems to outperform the frequency of antigen-specific PB (relative to the frequency of mBCs) to SARS-CoV-2 shortly after mRNA booster vaccination (25). In this case, the frequency of virus-specific PB equaled the number of mBCs at day 7 after booster, but considerably dropped rapidly thereafter. At day 14 after vaccination, PB were virtually absent. Likewise, vaccine-induced mBCs expressing a phenotype associated with recent activation (CD21^lo^CD11c^+^), dropped more abruptly than conventional mBCs. Similar findings have been described for influenza vaccination or infection (25, 26). These data are intriguing as they point to the notion that the constitution of the autoreactive B cell response resembles the composition of B cell responses to viral antigens shortly after vaccination. The autoreactive B cell response, however, does not transit to a resting state but remains continuously activated and able to form PBs without signs of exhaustion or decay. In this context, it relevant to note that the continuous activation of autoreactive responses can likely persist for years as some RA-patients had longstanding disease. This contrasts other situations, in which it is indicated that continuous antigenic triggering induces cellular exhaustion or decay (27, 28). Resistance of PTM-directed B cell responses to such mechanisms may be an important factor in the maintenance of disease chronicity. Likewise, all patients in our study had low to moderate disease activity at the time of sampling, indicating that autoreactive B cell responses maintain their state of activation despite the efficient therapeutic suppression of synovial inflammation. From a clinical point of view, this is perhaps not surprising, as only 10-15% of ACPA^+^ RA patients can successfully taper immunosuppressive medication over the course of their disease, while all others will eventually flare. It is tempting to speculate that the activated phenotype of the autoreactive B cell response contributes to flaring as our data clearly indicate sustained “immunological disease activity”, i.e. the absence of immunological remission despite clinical remission induced by treatment.

In chronic infections and AIDs with persistent antigen, DN2 B cells, which are CD21^-^, are often found to arise (29). In SLE, these B cells have been shown to produce autoantibodies (8). This subset has also been found to increase in RA and to associate with joint damage (30). We observed an enrichment in DN2 cells in some but not all patients. However, no enrichment of PTM-directed B cells in the DN2 population could be noted since the CD21^-^ PTM-directed B cells were CD27^+^. This indicates that the increase in DN2 is not directly related to autoantibody production. The notion that PTM-directed B cells are CD21^-^ is also enforced by the high expression of CXCR3, a chemokine receptor which is induced by T-bet (30, 31). Expression of CXCR3 allows for migration of cells to its ligands CXCL9, CXCL10 and CXCL11. As increased concentrations of these chemokines have been observed in synovial tissue, it is likely that the synovial microenvironment can attract PTM-directed B cells efficiently (32). Indeed, increased concentrations of ACPA and Cit-directed PB have been observed at high frequency in synovial fluid previously (14, 33).

Cit-, CarP and Acetyl-directed B cells were predominantly found in RA patients, while low frequencies of these B cells were found in HD as part of the resting naïve B cell population. This finding is in-line with recent indications attributing a protective function to ACPA and to anti-PTM responses in the context of e.g. infections. Notably, fluctuating AMPA reactivity has been described in first-degree relatives of patients with RA, while only the generation of a persisting, class-switched AMPA-response associated with disease (34). The latter raises the intriguing question as to the origin of the AMPA response. Cross-reactivity and the responsiveness to external antigens may be crucial in this context. Our study provides formal evidence that not only secreted AMPA, but also PTM-directed B cells are cross-reactive towards different PTMs. This is in line with data obtained from the study of monoclonal ACPA which were tested for the recognition of different PTMs in ELISA (12, 13, 35). Around 75% of these antibodies recognized another PTMs next to citrulline in ELISA. Interestingly, this percentage is considerably higher than the 20% observed in the present study. The difference could be explained by experimental set-up, as different PTM antigens coated separately in ELISA will also allow binding of monoclonal antibodies (mAb) that are of lower avidity to a specific PTM. The flow cytometry setting applied in this study, however, may more closely reflect the situation in-vivo in which differences in binding avidity and antigen density will generate competition. We observed that individual tetramers indeed reduced binding to one of the other PTMs (figure S5). While we may therefore underestimate the degree of cross-reactivity of the cellular AMPA-response, it is notable that we observed both cross-reactive and mono-specific cells. This is in-line with data from crystallization studies that indicate cross-reactive and ‘private’ recognition profiles of ACPA (36). We observed a higher percentage of cross-reactivity in the CarP and Acetyl-directed B cell population compared to the Cit-directed B cell population. These data, therefore, suggest that PTM-directed B cells likely have the highest affinity for citrullinated epitopes. Nonetheless, B cells will most likely be exposed to all PTMs in the body. As our data show that modified antigen-specific B cells are cross-reactive to different PTMs, these cells can become activated by a plethora of antigens if available in sufficient abundance. This may explain their continuous stage of activation, but also shows that multiple triggers, both internal or external, may drive these PTM-directed responses in individual patients.

Finally, the phenotype of PTM-directed B cells could provide a rationale to explore novel targets for the treatment of RA. Patients with highly active mBC and PB compartments, for example, could benefit from interventions that more specifically target both compartments. The expression of CD38 but also the high expression of CD19 may be useful in this respect given the recent advances made to deplete B cells by CD19-directed CAR T-cells in several rheumatic autoimmune diseases as well as the targeted depletion of PBs by anti-CD38 therapy (3, 37).

In conclusion, we here present a deep phenotyping approach to elucidate features of autoreactive B cells in RA. Our work revealed that these B cells, present as distinct cross-reactive populations within the larger B cell pool, have the phenotype of recently activated mBC and PB. They express high levels of CXCR3 yet lack markers associated with mucosal homing. In the context of RA, and possibly also in other AIDs, the long-term persistence of such activated cells may be one of the factors that determine chronicity of disease. Elucidation of the basic principles allowing the autoreactive B cells to continuously respond, replicate and differentiate without showing signs of exhaustion will likely be relevant to increase the understanding of human AIDs, but can also have implications for vaccination strategies and cancer immunotherapy.

## Methods

### Patients and healthy donors

Peripheral blood (45ml) was collected from 16 ACPA+ RA-patients visiting the outpatient clinic of the Department of Rheumatology at Leiden University Medical Center (LUMC). Peripheral blood of 9 healthy donors was collected via the LUMC voluntary donor service (LuVDS). All patients and healthy donors (HD) signed an IRB-approved informed consent form. Patient and HD characteristics are described in table S1.

### Antigen labeling and titration

Biotinylated CCP4 peptides (peptides of a fourth generation of cyclic citrullinated peptides (13), peptide sequences in table S2) were coupled to streptavidin-APC or streptavidin-BV605; biotinylated CHcitP4 was coupled to streptavidin-BV650 or streptavidin-BUV737, biotinylated CAcetylP4 was coupled to streptavidin-BV711 or streptavidin-BUV661, and the respective unmodified backbones CArgP4 and CLysP4 were coupled to streptavidin-APC-Fire750. All coupling steps were performed by incubation overnight at 4°C. The next day, free peptides were removed using a Bio-Spin P-30 column (Biorad), leaving the labeled PTM tetramers in the flowthrough. Tetanus Toxoid (TT) was labeled with APC or PE using an AnaTag labeling kit (Anaspec) according to the manufacturer’s protocol. The labeled antigens were titrated on MDL-AID KO Ramos cells transduced with BCRs reactive to PTM-antigens (13). For CCP4 and CHcitP4, the 2G9 cell line was used and for CAcetylP4, the 7E4 cell line was used. Labeled TT was titrated on immortalized TT-specific B cells, as previously described (14).

### Flow cytometry

PBMCs were isolated using Ficoll-plaque gradient centrifugation and stored overnight at 4°C in RPMI+8%FCS+glutamax+penicillin/streptomycin. The next day, B cells were enriched using Easysep B cell purification kit (Stemcell) following the manufacturer’s protocol. B cells were then stained with Fixable Viability Dye for 30 min on ice in the dark. After washing twice with PBS, cells were stained with CXCR3, CCR7 and CCR9 and incubated at 37°C in the dark for 30 min. Next, cells were washed twice with PBS+1%BSA and stained with CD3, CD14, CD19, CD20, CD21, CD24, CD27, CD38, CD62L, CD80, CD86, IgD and fluorochrome-labeled antigens and incubated for 30 min on ice in the dark. After washing twice with PBS+1%BSA, cells were fixed and permeabilized with FoxP3/Transcription factor staining buffer set (eBioscience). Cells were subsequently stained with IgM, IgA, IgG, Ki-67 and the fluorochrome-labeled antigens and incubated for 30 min on ice in the dark. After washing twice with permeabilization buffer, the cells were resuspended in PBS+1%BSA and measured with a 5-laser Cytek Aurora flow cytometer. Reference PBMCs, pooled buffy coats from 2 healthy donors, were taken along in every experiment to adjust for variation in fluorochrome intensity in different experiments, resulting in a ‘normalized MFI’. The fluorochromes, clones, category numbers and dilutions of the antibodies can be found in table S3.

### Data analysis

Flow cytometry data were analyzed using the OMIQ software from Dotmatics (www.omiq.ai, www.dotmatics.com). Antigen-specific B cells were gated manually per experiment. Data from different experiments were normalized with Cytonorm using pooled PBMCs from 2 healthy donor buffy coats (38). Then, unsupervised FlowSom clustering was performed using normalized CD19, CD20, CD21, CD24, CD27, CD38, IgD and IgM to identify different B cell clusters (16, 39). Data were exported from OMIQ and visualization and statistical testing was performed using Graphpad Prism 9.3.1. Samples with less than 10 cells per group were excluded from the analysis. Differences in frequencies of antigen-specific B cells or B cell subsets between HD and RA patients were statistically tested with Mann-Whitney U tests. Differences in marker expression and B cell percentages in different B cell clusters were statistically tested with the Wilcoxon signed rank test.

## Author Contributions

SR and JK performed experiments; NJB, JS, SR designed panel; DW, REMT and HUS supervised the project. SR analyzed the data and drafted the manuscript. All authors provided feedback and approved submission of the manuscript.

## Acknowledgments

The authors would like to thank Jan Wouter Drijfhout for synthesizing the peptides and the Flow Cytometry Core Facility of the LUMC for assistance with the spectral analyzers. We thank Sophie-Anne Smith, MSc for help in preparing figures.

## References

1. Edwards JCW, Szczepanski L, Szechinski J, et al. Efficacy of B-Cell–Targeted Therapy with Rituximab in Patients with Rheumatoid Arthritis. New England Journal of Medicine. 2004;350(25):2572–81.

2. Joly P, Maho-Vaillant M, Prost-Squarcioni C, et al. First-line rituximab combined with short-term prednisone versus prednisone alone for the treatment of pemphigus (Ritux 3): a prospective, multicentre, parallel-group, open-label randomised trial. The Lancet. 2017;389(10083):2031–40.

3. Mackensen A, Müller F, Mougiakakos D, et al. Anti-CD19 CAR T cell therapy for refractory systemic lupus erythematosus. Nature Medicine. 2022;28(10):2124–32.

4. Young C, and McGill SC. CADTH Health Technology Review. 2021.

5. Amanna IJ, Carlson NE, and Slifka MK. Duration of humoral immunity to common viral and vaccine antigens. N Engl J Med. 2007;357(19):1903–15.

6. Phad GE, Pinto D, Foglierini M, et al. Clonal structure, stability and dynamics of human memory B cells and circulating plasmablasts. Nat Immunol. 2022;23(7):1–10.

7. Moir S, Buckner CM, Ho J, et al. B cells in early and chronic HIV infection: evidence for preservation of immune function associated with early initiation of antiretroviral therapy. Blood. 2010;116(25):5571–9.

8. Wang S, Wang J, Kumar V, et al. IL-21 drives expansion and plasma cell differentiation of autoreactive CD11c(hi)T-bet(+) B cells in SLE. Nat Commun. 2018;9(1):1758.

9. Qin Y, Cai ML, Jin HZ, et al. Age-associated B cells contribute to the pathogenesis of rheumatoid arthritis by inducing activation of fibroblast-like synoviocytes via TNF-α-mediated ERK1/2 and JAK-STAT1 pathways. Ann Rheum Dis. 2022;81(11):1504–14.

10. Reed E, Jiang X, Kharlamova N, et al. Antibodies to carbamylated alpha-enolase epitopes in rheumatoid arthritis also bind citrullinated epitopes and are largely indistinct from anti-citrullinated protein antibodies. Arthritis Res Ther. 2016;18(1):96.

11. Kampstra ASB, Dekkers JS, Volkov M, et al. Different classes of anti-modified protein antibodies are induced on exposure to antigens expressing only one type of modification. Annals of the Rheumatic Diseases. 2019;78(7):908–16.

12. Sahlström P, Hansson M, Steen J, et al. Different Hierarchies of Anti–Modified Protein Autoantibody Reactivities in Rheumatoid Arthritis. Arthritis & Rheumatology. 2020;72(10):1643–57.

13. Kissel T, Reijm S, Slot L, et al. Antibodies and B cells recognising citrullinated proteins display a broad cross-reactivity towards other post-translational modifications. Annals of the Rheumatic Diseases. 2020:annrheumdis-201.

14. Kristyanto H, Blomberg NJ, Slot LM, et al. Persistently activated, proliferative memory autoreactive B cells promote inflammation in rheumatoid arthritis. Science Translational Medicine. 2020;12(570):eaaz5327.

15. Kerkman PF, Fabre E, van der Voort EIH, et al. Identification and characterisation of citrullinated antigen-specific B cells in peripheral blood of patients with rheumatoid arthritis. Annals of the Rheumatic Diseases. 2016;75(6):1170–6.

16. Sanz I, Wei C, Jenks SA, et al. Challenges and Opportunities for Consistent Classification of Human B Cell and Plasma Cell Populations. Front Immunol. 2019;10:2458.

17. Scharer CD, Patterson DG, Mi T, et al. Antibody-secreting cell destiny emerges during the initial stages of B-cell activation. Nature Communications. 2020;11(1).

18. Macpherson AJ, McCoy KD, Johansen FE, et al. The immune geography of IgA induction and function. Mucosal Immunol. 2008;1(1):11–22.

19. Mora JR, Iwata M, Eksteen B, et al. Generation of gut-homing IgA-secreting B cells by intestinal dendritic cells. Science. 2006;314(5802):1157–60.

20. Muehlinghaus G, Cigliano L, Huehn S, et al. Regulation of CXCR3 and CXCR4 expression during terminal differentiation of memory B cells into plasma cells. Blood. 2005;105(10):3965–71.

21. Baumgarth N. The Shaping of a B Cell Pool Maximally Responsive to Infections. Annu Rev Immunol. 2021;39:103–29.

22. Reincke ME, Payne KJ, Harder I, et al. The Antigen Presenting Potential of CD21(low) B Cells. Front Immunol. 2020;11:535784.

23. Lau D, Lan LY, Andrews SF, et al. Low CD21 expression defines a population of recent germinal center graduates primed for plasma cell differentiation. Sci Immunol. 2017;2(7).

24. Sokal A, Chappert P, Barba-Spaeth G, et al. Maturation and persistence of the anti-SARS-CoV-2 memory B cell response. Cell. 2021;184(5):1201-13.e14.

25. Kardava L, Rachmaninoff N, Lau WW, et al. Early human B cell signatures of the primary antibody response to mRNA vaccination. Proc Natl Acad Sci U S A. 2022;119(28):e2204607119.

26. Ellebedy AH, Jackson KJ, Kissick HT, et al. Defining antigen-specific plasmablast and memory B cell subsets in human blood after viral infection or vaccination. Nat Immunol. 2016;17(10):1226–34.

27. Shimabukuro-Vornhagen A, García-Márquez M, Fischer RN, et al. Antigen-presenting human B cells are expanded in inflammatory conditions. J Leukoc Biol. 2017;101(2):577–87.

28. Moir S, Ho J, Malaspina A, et al. Evidence for HIV-associated B cell exhaustion in a dysfunctional memory B cell compartment in HIV-infected viremic individuals. J Exp Med. 2008;205(8):1797–805.

29. Cooper L, and Good-Jacobson KL. Dysregulation of humoral immunity in chronic infection. Immunol Cell Biol. 2020;98(6):456–66.

30. Thorarinsdottir K, Camponeschi A, Jonsson C, et al. CD21(-/low) B cells associate with joint damage in rheumatoid arthritis patients. Scand J Immunol. 2019;90(2):e12792.

31. Ly A, Liao Y, Pietrzak H, et al. Transcription Factor T-bet in B Cells Modulates Germinal Center Polarization and Antibody Affinity Maturation in Response to Malaria. Cell Reports. 2019;29(8):2257-69.e6.

32. Ueno A, Yamamura M, Iwahashi M, et al. The production of CXCR3-agonistic chemokines by synovial fibroblasts from patients with rheumatoid arthritis. Rheumatol Int. 2005;25(5):361–7.

33. Willemze A, Shi J, Mulder M, et al. The concentration of anticitrullinated protein antibodies in serum and synovial fluid in relation to total immunoglobulin concentrations. Annals of the Rheumatic Diseases. 2013;72(6):1059–63.

34. Tanner S, Dufault B, Smolik I, et al. A Prospective Study of the Development of Inflammatory Arthritis in the Family Members of Indigenous North American People With Rheumatoid Arthritis. Arthritis Rheumatol. 2019;71(9):1494–503.

35. Reijm S, Kissel T, Stoeken-Rijsbergen G, et al. Cross-reactivity of IgM anti-modified protein antibodies in rheumatoid arthritis despite limited mutational load. Arthritis Research & Therapy. 2021;23(1).

36. Ge C, Xu B, Liang B, et al. Structural Basis of Cross-Reactivity of Anti-Citrullinated Protein Antibodies. Arthritis Rheumatol. 2019;71(2):210–21.

37. Ostendorf L, Burns M, Durek P, et al. Targeting CD38 with Daratumumab in Refractory Systemic Lupus Erythematosus. N Engl J Med. 2020;383(12):1149–55.

38. Van Gassen S, Gaudilliere B, Angst MS, et al. CytoNorm: A Normalization Algorithm for Cytometry Data. Cytometry Part A. 2020;97(3):268–78.

39. Van Gassen S, Callebaut B, Van Helden MJ, et al. FlowSOM: Using self-organizing maps for visualization and interpretation of cytometry data. Cytometry A. 2015;87(7):636–45.

