## Supplemental file for "Autoreactive B cells in rheumatoid arthritis consist of activated CXCR3+ memory B cells and plasmablasts"

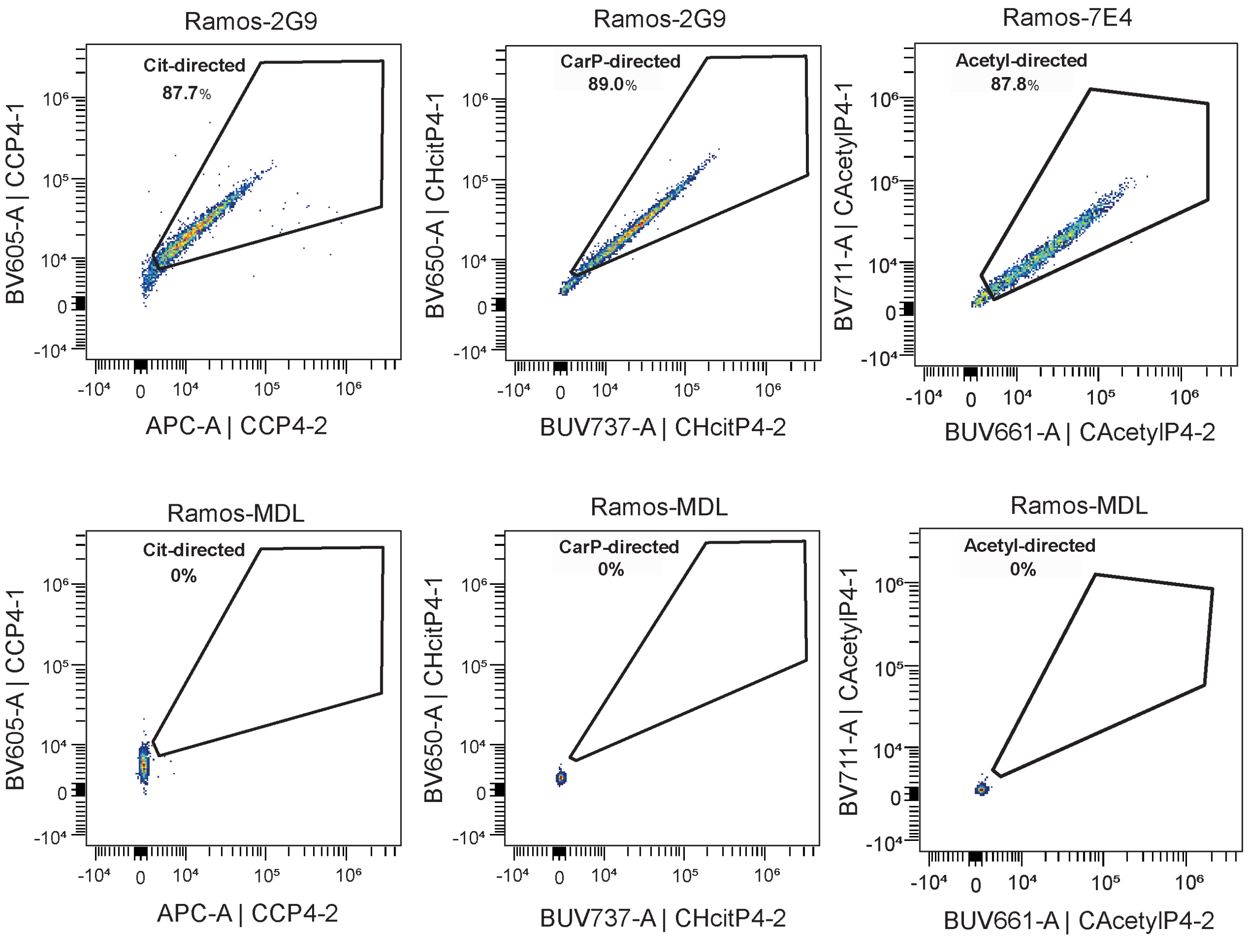


Figure S1. Tetramer staining on Ramos-2G9 or Ramos-7E4 (upper panel) or Ramos-MDL cell lines (lower panel). Cit-directed B cell staining on the left, CarP-directed B cell staining in the middle and Acetyl-directed B cell staining on the right. N = 5, one representative experiment is shown in the figure.

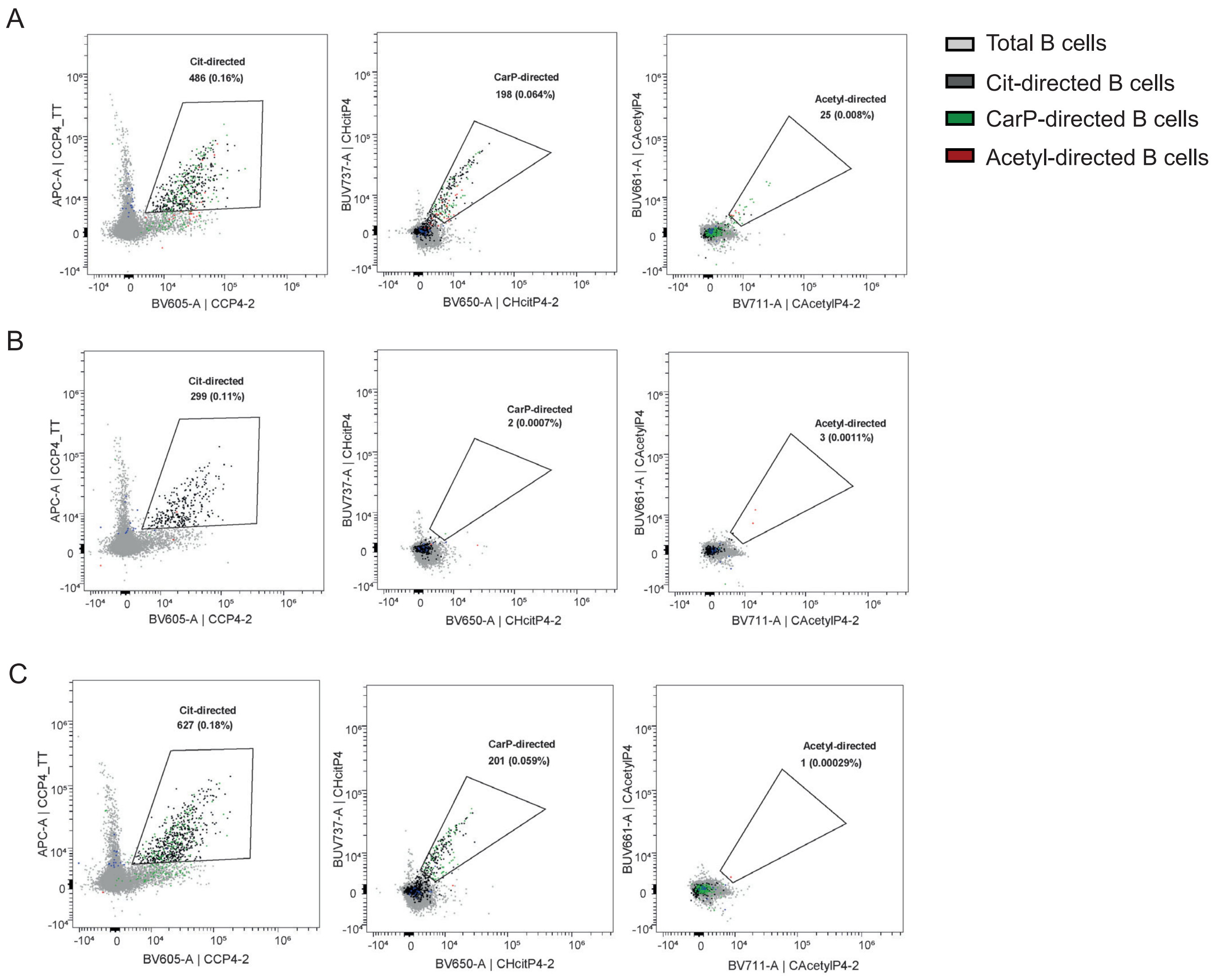


Figure S2. Blocking experiments. Cit-directed B cell staining (left), CarP-directed B cell staining (middle) and Acetyl-directed B cell staining (right) of cells from one RA patient. In A, the staining was not inhibited. In B, the signal was blocked with unlabeled CHcitP4 tetramers before staining with the labeled tetramers. In C, the signal was blocked with unlabeled CAcetylP4 tetramers before staining with the labeled tetramers.

A

B

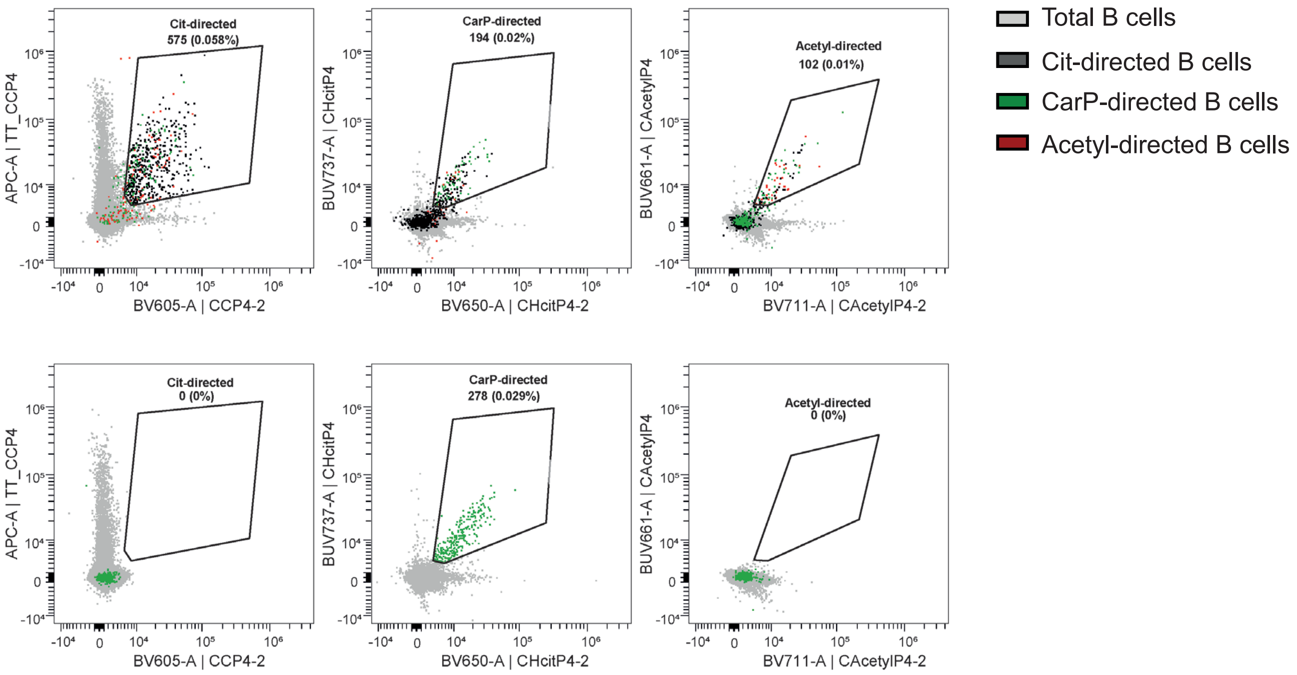


Figure S3. Cit-directed B cell staining (left), CarP-directed B cell staining (middle) and Acetyl-directed B cell staining (right) of cells from one RA patient. In A, labeled tetramers for all PTMs were added. In B, only labeled CHCitP4 tetramers were added.

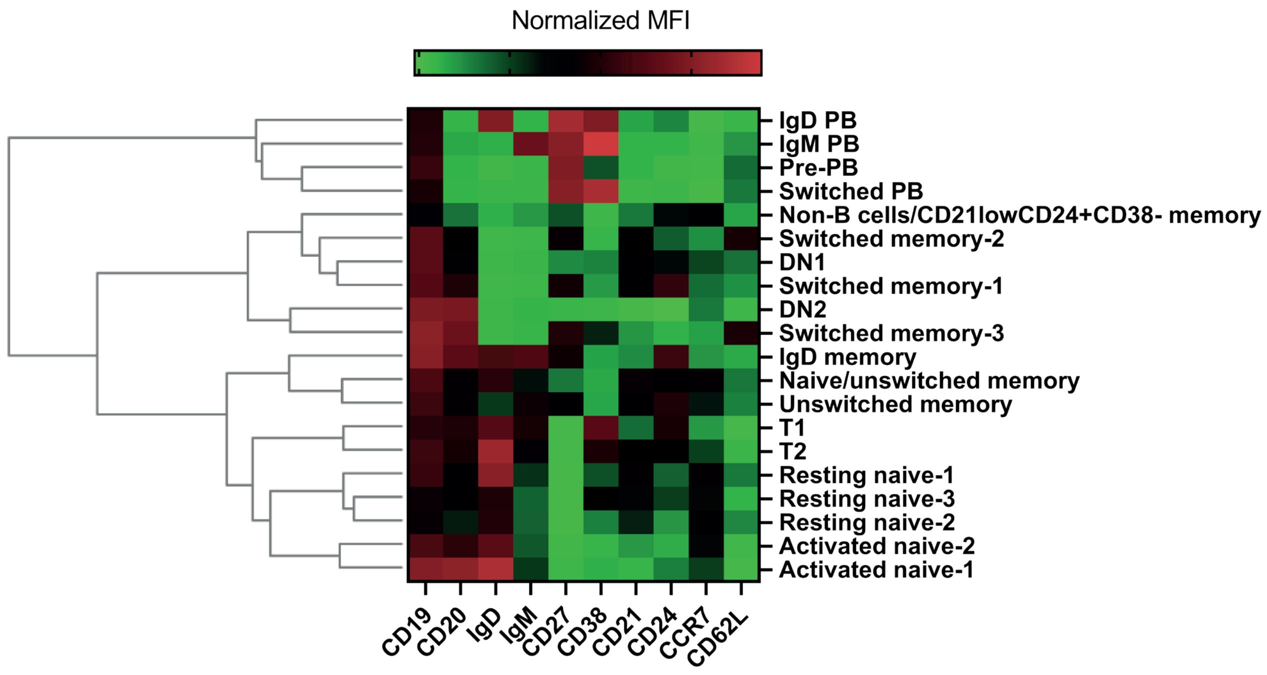


Figure S4. Heatmap with hierarchical clustering of the 20 B cell clusters obtained using the clustered heatmap function in OMIQ. From all samples, 30k B cells were subsampled and the normalized MFI (green is low, red is high MFI) was used for for this analysis.

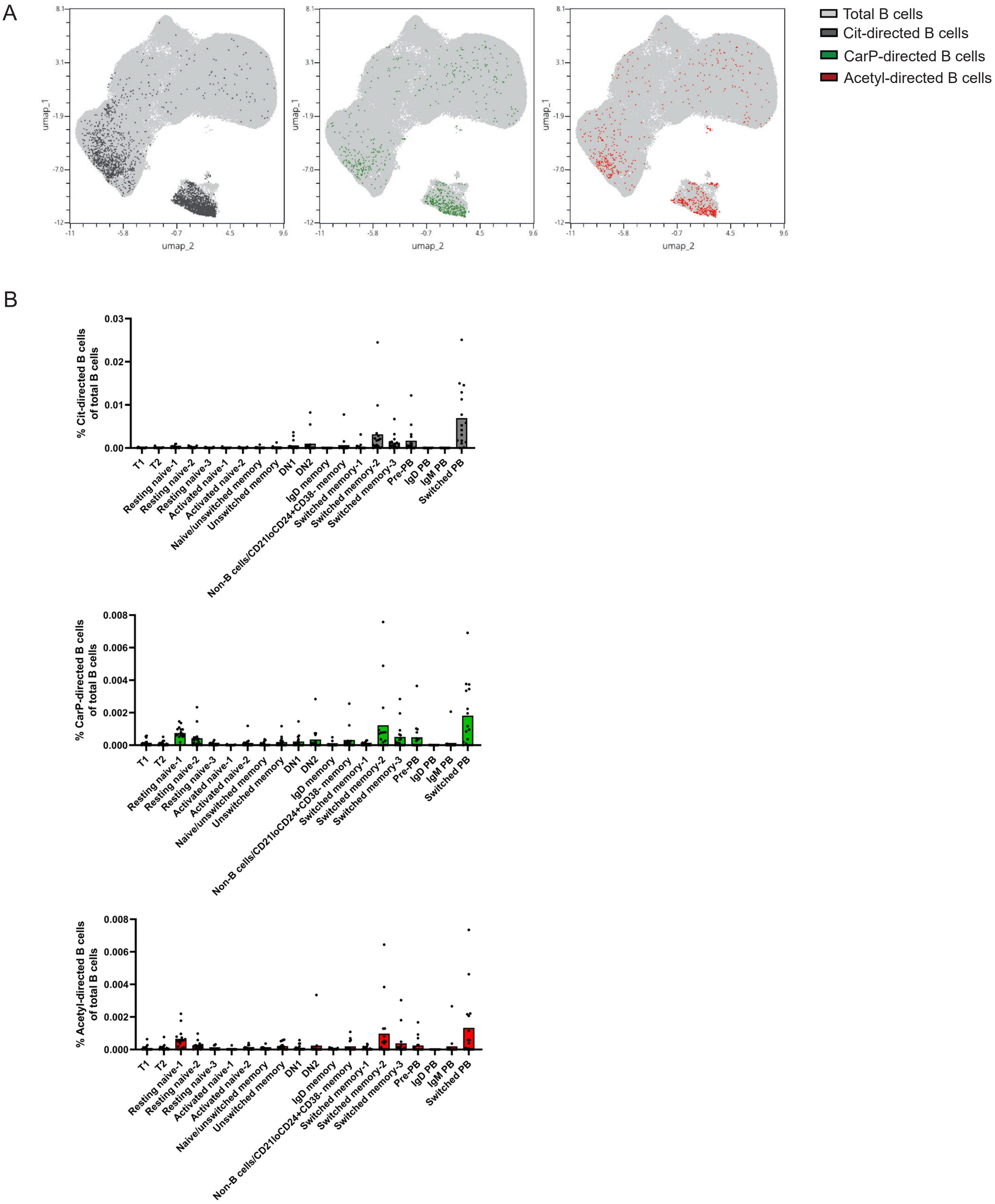


Figure S5. A) UMAP visualization of total B cells (light grey), Cit-directed B cells (dark grey), CarP-directed B cells (green) and Acetyl-directed B cells (red) of RA patients (n=16). B) Percentage of Cit-, CarP- and Acetyl-directed B cells of total B cells separated by the 20 identified B cell subsets. Each dot represents one individual, bars represent means.

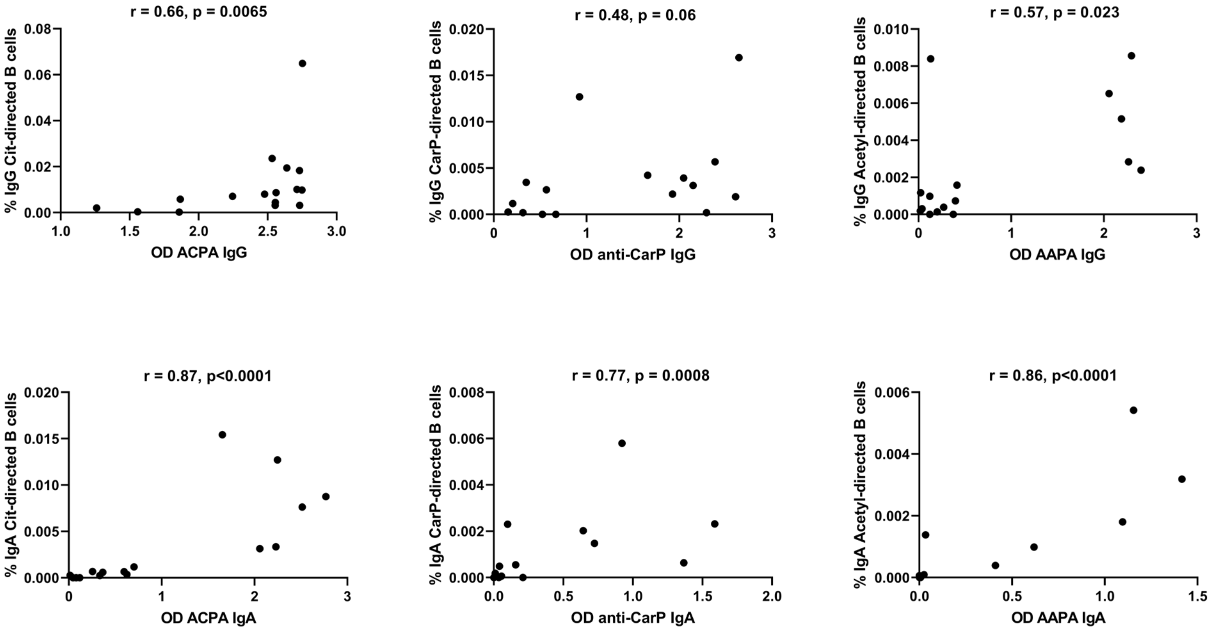


Figure S6. Correlation of the percentage of Cit-, CarP or Acetyl-directed cells of total B cells with plasma AMPA levels for IgG (upper panel) and IgA (lower panel).

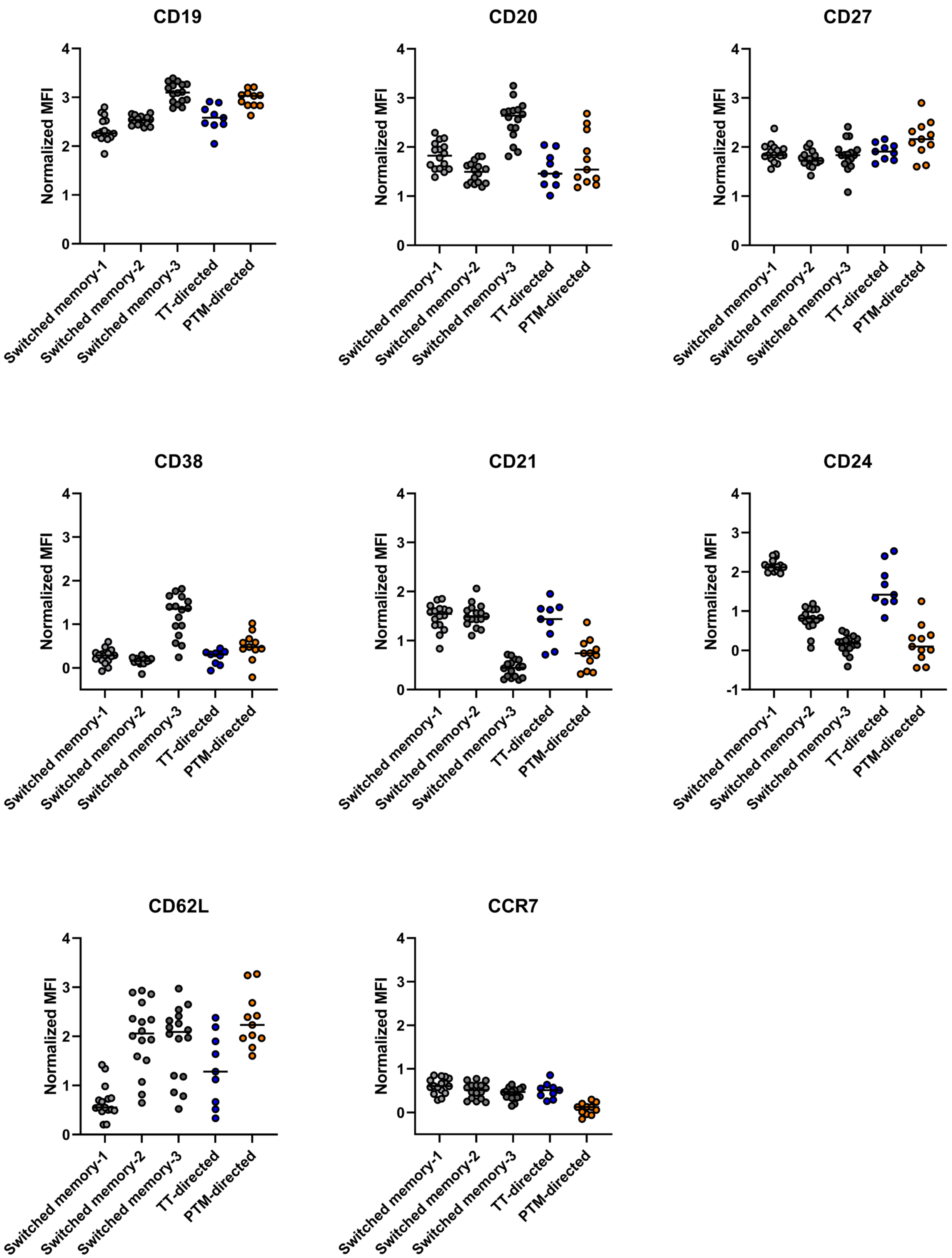


Figure S7. Normalized MFI of CD19, CD20, CD27, CD38, CD21 CD24, CCR and CD62L of 3 switched memory clusters and TT-directed B cells and PTM-directed B cells within these clusters. Each dot represent 1 RA patient. Only the PTM- and TT-directed B cells that were present in 1 of the 3 switched memory clusters were included in the analysis. Donors with less than 10 TT- or PTM-directed B cells in the analyzed clusters were excluded from the analysis.

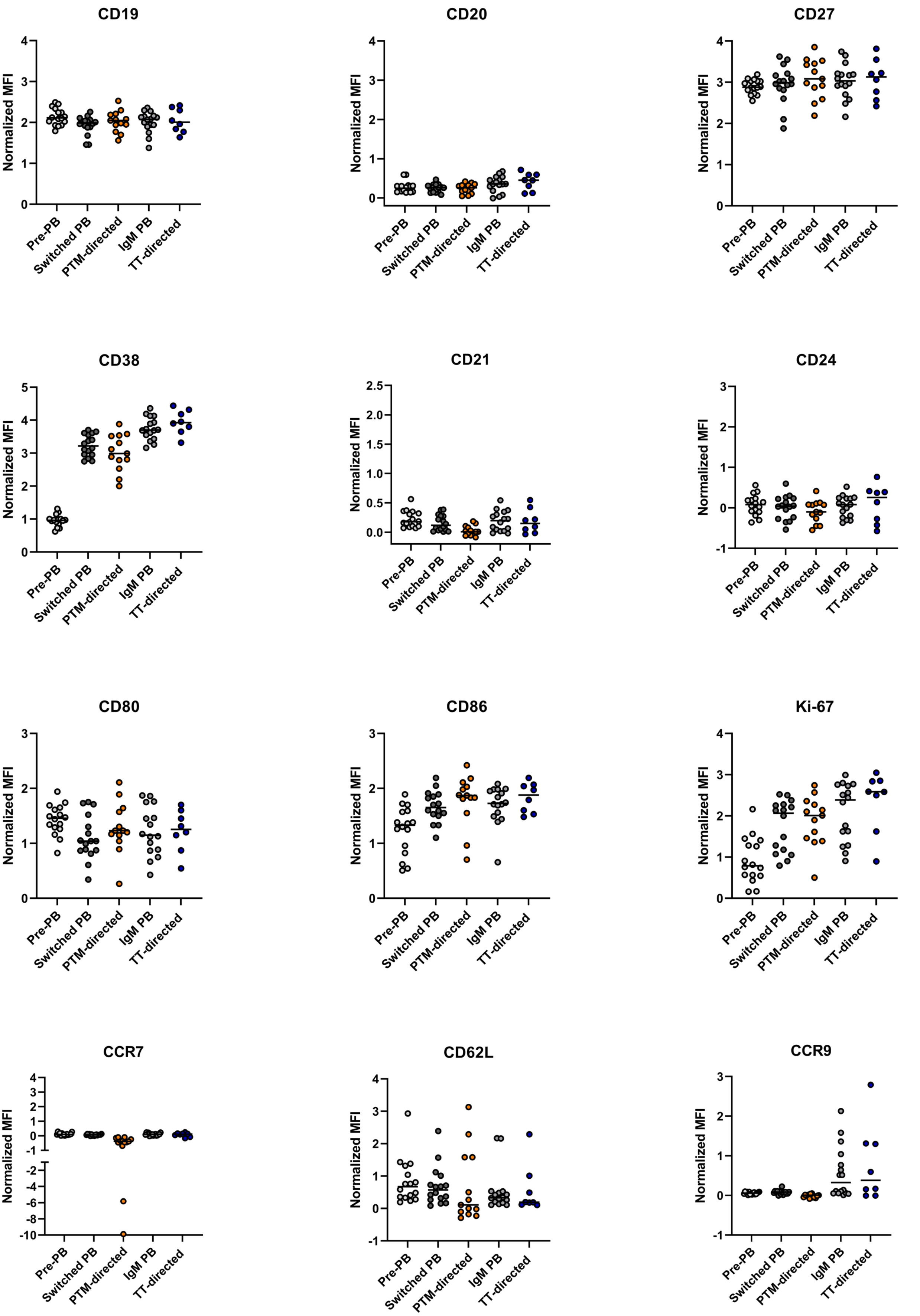


Figure S8. Normalized MFI of CD19, CD20, CD27, CD38, CD21, CD24, CD80, CD86, Ki-67, CCR7, CD62L and CCR9 of 3 PB clusters and TT-directed B cells and PTM-directed B cells within these clusters. Each dot represent 1 RA patient. Only the AMPA- and TT-directed B cells that were present in 1 of the 3 PB clusters were included in the analysis. Donors with less than 10 TT- or PTM-directed B cells in the analyzed clusters were excluded from the analysis.

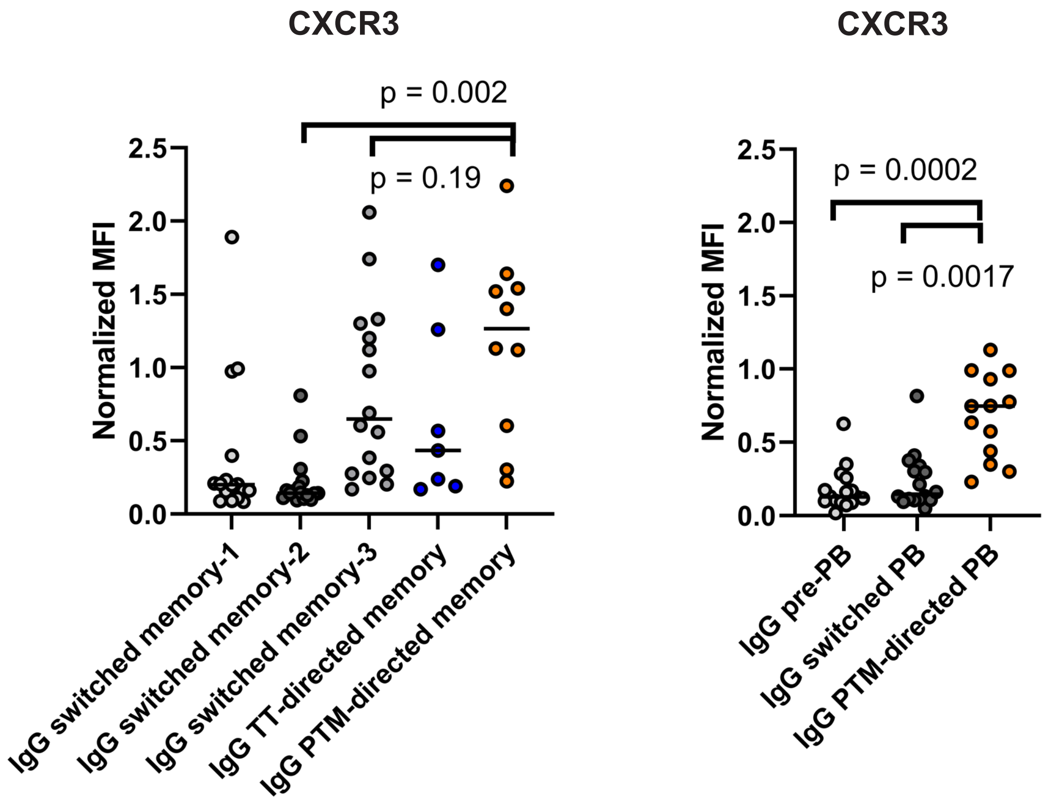


Figure S9. Normalized MFI of CXCR3 of IgG+ cells in the switched memory clusters and TT-directed B cells and PTM-directed B cells within these clusters (right). Normalized MFI of CXCR3 of IgG+ cells in the PB clusters and PTM-directed B cells within these clusters (left). Statistical testing was performed with Wilcoxon signed rank test. Each dot represent 1 RA patient. Only the PTM- and TT-directed B cells that were present in 1 of the memory or PB clusters were included in the analysis. Donors with less than 10 TT- or PTM-directed B cells in the analyzed clusters were excluded from the analysis.

**Table S1 patient characteristics**

| | RA patients (16) | Healthy donors (9) |
|---|---|---|
| **Sex (%)** | | |
| Male | 25 | 22.2 |
| Female | 75 | 77.8 |
| **Age (years)** | | |
| Median | 67.5 | 54 |
| Range | 42-87 | 36-66 |
| **Most recent CCP titer (U/ml)** | | |
| Median | >300 | n.a. |
| Range | 287-600 | n.a. |
| **DAS28** | | |
| Median | 2.69 | n.a. |
| Range | 0.68-4.01 | n.a. |
| **Treatment at the time of sampling (n)** | | |
| Methotrexate | 9 | n.a. |
| Sulfasalazine | 2 | n.a. |
| Leflunomide | 1 | n.a. |
| Anti-IL-6 | 1 | n.a. |
| Anti-TNFα | 2 | n.a. |
| Prednisone | 2 | n.a. |
| Etoricoxib | 1 | n.a. |

**Table S2 Peptide sequences**

| Peptide | Sequence |
|---|---|
| CCP4 (cyclic) | HQFRFXGNleSRAACZO |
| CArgP4 (cyclic) | HQFRFRGNleSRAACZO |
| CHcitP4 (cyclic) | HQFRFJGNleSRAACZO |
| CAcetylP4 (cyclic) | HQFRFBGNleSRAACZO |
| CLysP4 (cyclic) | HQFRFKGNleSRAACZO |

X = citrulline
J = homocitrulline
B = acetyllysine
Z = 6-aminohexanoic acid
O = lys (biotine)-amide
Nle= norleucine

**Table S3. Fluorochromes, dilutions, clones and catalog numbers of the antibodies**

| Antibody | Dilution | Clone | Company | Catalog |
|---|---|---|---|---|
| Fixable Viability Dye eFluor506 | 1000 | n.a. | eBioscience | 65-0866-14 |
| CD3-eFuor506 | 25 | UCHT1 | eBioscience | 69-0038-42 |
| CD14-efluor506 | 25 | 61D3 | eBioscience | 69-0149-42 |
| CD19-BV570 | 25 | HIB19 | Biolegend | 302236 |
| CD20-BUV395 | 50 | 2H7 | BD biosciences | 563782 |
| CD21-BUV805 | 400 | B-ly4 | BD biosciences | 742008 |
| CD24-BUV496 | 25 | ML5 | BD biosciences | 741143 |
| CD27-APCFire810 | 100 | QA17A18 | Biolegend | 393214 |
| CD38-BUV563 | 1000 | HB7 | BD biosciences | 741446 |
| CD62L-SparkNIR685 | 400 | DREG-56 | Biolegend | 304861 |
| CD80-BB515 | 400 | L307.4 | BD bioscience | 565008 |
| CD86-BV785 | 50 | IT2.2 | Biolegend | 305441 |
| CXCR3/CD183-BB700 | 50 | 1C6 | BD bioscience | 566533 |
| CCR7/CD197-Pe-Fire640 | 100 | G043H7 | Biolegend | 353261 |
| CCR9/CD199 -Pe-cy7 | 800 | L053E8 | Biolegend | 358909 |
| IgD-BV480 | 400 | IA6-2 | BD biosciences | 566138 |
| IgA-VioBlue | 400 | IS11-8E10 | Miltenyi | 130-113-479 |
| IgG-BV421 | 50 | G18-145 | BD biosciences | 562581 |
| IgM-SB436 | 50 | SA-DA4 | eBioscience | 62-9998-42 |
| Ki-67-PE-eFluor610 | 2000 | 20Raj1 | eBioscience | 61-5699-41 |
| Streptavidin-APC | n.a. | n.a. | Invitrogen | S32362 |
| Streptavidin-APC-Fire750 | n.a. | n.a. | Biolegend | 405250 |
| Streptavidin-BUV661 | n.a. | n.a. | BD bioscience | 612979 |
| Streptavidin-BUV737 | n.a. | n.a. | BD bioscience | 612775 |
| Streptavidin-BV605 | n.a. | n.a. | Biolegend | 405229 |
| Streptavidin-BV650 | n.a. | n.a. | Biolegend | 405231 |
| Streptavidin-BV711 | n.a. | n.a. | Biolegend | 405241 |
